# Tension shapes memory: Computational insights into neural plasticity

**DOI:** 10.1101/2025.08.20.671220

**Authors:** Ki Yun Lee, M. Taher A. Saif

**Affiliations:** Department of Mechanical Science and Engineering, University of Illinois at Urbana-Champaign, Urbana, IL 61801, USA; Beckman Institute for Advanced Science and Technology, University of Illinois at Urbana-Champaign, Urbana, IL 61801, USA

**Author notes:** Corresponding authors (MS).

## Abstract

Mechanical forces have recently emerged as critical modulators of neural communication, yet their role in high-level cognitive functions remains poorly understood. Here, we present a biologically inspired spiking neural network model that integrates mechanical tension, vesicle dynamics, and spike-timing-dependent plasticity to examine how tension influences learning, memory, and cognitive operations such as pattern completion, projection, and association. We find that increased tension enhances synaptic efficiency by accelerating vesicle clustering and recovery, resulting in a 67% improvement in memory recall speed and a 17% increase in inter-regional synchrony during projection relative to relaxed states. Conversely, a 20% reduction in tension leads to a 31% decline in memory association performance, highlighting the tension-sensitive accessibility of stored information. The model further reveals that networks with 20% inhibitory neurons achieve optimal spatial precision in memory encoding and recall. Together, these *in silico* findings position mechanical tension as a functional neuromodulator and suggest new directions for neuromorphic design and energy-efficient, living computing platforms.

**Author summary:** Our brains are not only electrical and chemical systems but also mechanical ones. Neurons and their connections are constantly under tension, yet the role of these physical forces in shaping memory and thought remains poorly understood. In this work, we built a computer model of a brain-like network that allowed us to test how mechanical tension influences learning and memory. We found that higher tension improved the speed and accuracy of memory recall, strengthened communication between groups of neurons, and supported more reliable association of memories. In contrast, reducing tension made memories harder to access, though they could be recovered once tension was restored. We also discovered that having the right balance between excitatory and inhibitory neurons was crucial for both storing and recalling information. Our results suggest that tension is not just a background property of brain tissue but an active player in cognition, with potential implications for understanding memory disorders and designing new brain-inspired technologies.

## Introduction

Neural communication is fundamental to high-level cognitive functions such as abstract thinking, decision-making, learning, and memory. Traditionally, these processes have been studied through chemical signaling mechanisms such as ion flux, neurotransmitter release, and receptor binding [1,2]. However, growing evidence suggests that mechanical forces, particularly tension arising from contractility of axons, also play a vital role in shaping neuronal behavior. The role of neuronal tension has been tested and verified by a wide range of experiments, both *in vitro* and *in vivo*. For instance, external forces can induce cytoskeletal remodeling and axonal outgrowth, processes central to neural plasticity [3–5]. Neurons generate and maintain contractile forces via the actomyosin cytoskeleton [6–8], and that tension increases during axon elongation and synapse formation. Notably, neurons may fail to function properly in the absence of tension, highlighting its importance not only for development, but also for ongoing neural activity, learning, and memory maintenance [9].

At the synaptic level, mechanical tension enhances vesicle clustering at presynaptic terminals, increases neural excitability, and promotes activity-dependent plasticity. Experimental works with *Drosophila* embryos demonstrated that disrupting axonal tension reduced vesicle accumulation, while restoring tension recovered it [10], and increasing tension further enhanced vesicle clustering [11] (Fig 1A). *Ex vivo* mouse brain slices under cyclic stretch exhibited cumulative increases in excitability, suggesting the presence of a “mechanical memory” [12]. Further, chemically modulating neural contractility altered vesicle accumulation and firing rates, reinforcing the idea that mechanical forces actively regulate neural signaling [9,13]. These findings collectively position tension not as a passive byproduct of cellular architecture but as an active neuromodulatory signal that supports plasticity, network synchronization, and memory-like behavior. Motivated by this, we developed a computational model of a spiking neural network under mechanical tension, incorporating excitatory and inhibitory leaky integrate-and-fire neurons, vesicle dynamics, and synaptic plasticity. We hypothesize that mechanical tension enhances synaptic strength and facilitates cognitive functions by increasing vesicle availability and modulating network dynamics.

**Fig 1.**
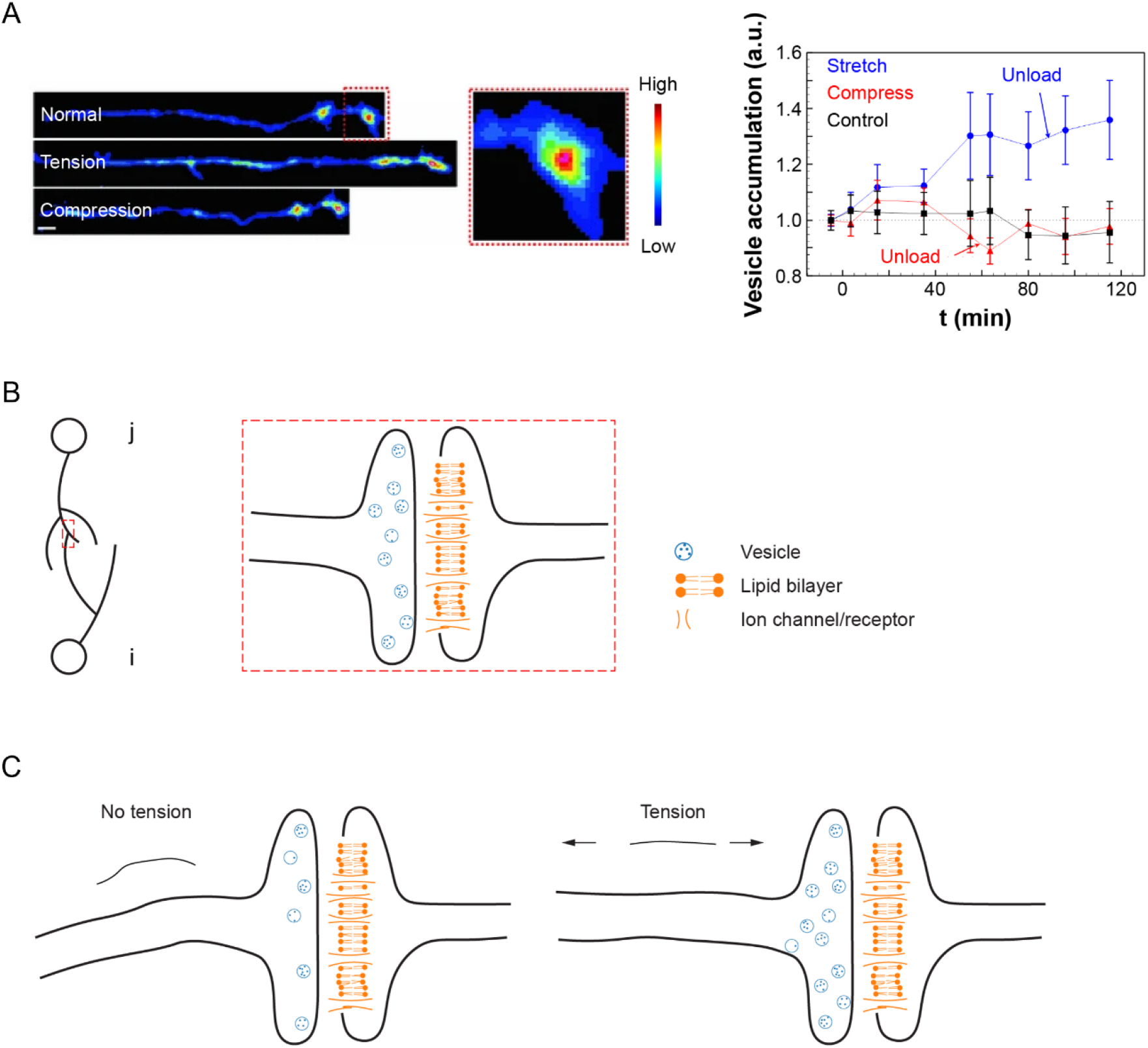
Experimental evidence for vesicle clustering and schematic illustration of the spiking neural network model under mechanical tension. (A) Drosophila embryo motor neurons (left) and synaptic vesicle accumulation at neuromuscular junctions (right; mean ± SEM) under different mechanical conditions, adapted from [11]. Vesicle accumulation increased by 30% under tension and remained elevated for at least 30 minutes after unloading (*n* = 8). Compressed axons showed no significant change over time (*n* = 8), and control samples also remained stable (*n* = 10). Scale bar: 5 µm. (B) Schematic of a synapse formed between presynaptic neuron *j* (containing neurotransmitter vesicles) and postsynaptic neuron *i* (with receptors at the postsynaptic membrane). (C) Illustration showing increased vesicle accumulation at the presynaptic terminal as mechanical tension increases.

A neural assembly refers to a group of interconnected neurons that tend to activate together within a network, forming functionally meaningful units [14–16]. According to Hebbian theory, neurons within an assembly respond collectively to stimuli rather than acting independently [16]. To model such behavior at a higher level of abstraction, a framework known as assembly calculus was developed [17]. This framework enables simulation of cognitive operations such as pattern completion (retrieving a full pattern from partial input), projection (forming a new assembly via activation of an existing one), and association (linking two assemblies such that activation of one triggers the other). Assembly calculus introduces control over network activation using parameters such as the total number of neurons (*n*) and the maximum number of simultaneously spiking neurons (*k*), with *k* effectively simulating inhibition by restricting widespread activation. While this symbolic approach enables computational modeling of high-level operations, it abstracts away the underlying biophysical mechanisms. In contrast, our model implements these cognitive functions within a biophysically realistic framework, capturing the dynamics of individual neurons, synapses, and vesicle clustering. This allows us to examine how cognitive operations might emerge from low-level, biologically grounded interactions in real neural tissue.

Hebbian theory posits that learning and memory result from activity-dependent changes in synaptic plasticity, such as synaptogenesis and adjustments in synaptic weight [16]. Repeated stimulation (i.e., learning) leads to the formation of assemblies that temporarily operate as closed systems. Once stabilized, these assemblies can function independently and support cognitive operations [14]. A key limitation of Hebbian learning is the unbounded growth of synaptic weights, which can destabilize the network by causing assemblies to merge into the broader network. Oja’s rule addresses this by introducing a decay term to stabilize synaptic weights [18]. In our model, we extend this principle with a biologically motivated forgetting mechanism supported by experimental evidence [19] and incorporate mechanical tension as a dynamic variable to probe its role as a neuromodulator. Using a biologically inspired spiking neural network composed of excitatory and inhibitory neurons, and integrating learning-based synaptic plasticity with vesicle dynamics, we examined how training duration and tension level, and inhibition influence pattern completion, memory encoding and recall, and the emergence of higher-order cognitive operations such as projection and association. Our simulations demonstrate successful pattern completion, projection, and associative memory recall following training, indicating functional plasticity in the trained network. These results suggest that mechanical tension shapes the dynamics of neural assemblies, influencing not only firing and plasticity but also inter-assembly interactions.

## Materials and Methods

### Spiking neural network model

We constructed a spiking neural network comprising 4,000 excitatory and 1,000 inhibitory neurons randomly distributed within a bounded 2D space. The 4:1 excitatory-to-inhibitory ratio was chosen based on physiological data from the rodent hippocampus [20,21] and primate cortex [22,23], as well as prior simulation studies [24,25]. Each neuron was assigned multiple neurites (dendrites and axons), with lengths drawn from a normal distribution (mean = 200 μm; standard deviation = 40 μm). Synapses were established probabilistically according to the Euclidean distance between neurons, with a fixed connection probability of 0.1. Self-connections were not permitted, and each presynaptic neuron could form at most one synapse with any given postsynaptic neuron.

### Neural communication dynamics

We modeled synaptic transmission between a presynaptic neuron *j* and a postsynaptic neuron *i* (Fig 1B). The presynaptic neuron clusters neurotransmitter-filled vesicles at its terminal, while the postsynaptic neuron clusters neurotransmitter receptors at the synaptic membrane. The membrane potential of neuron *i* at time *t*, denoted as *V*_*i*_*(t)*, is initialized at a resting potential *V*_*rest*_ in the absence of stimulation.

When an action potential (AP) arrives at the presynaptic terminal of neuron *j*, neurotransmitters are released and they bind to receptors on neuron *i*, inducing ion flux and increasing *V*_*i*_*(t)* by an amount *V*_*synapse*_*(t)*. If *V*_*i*_*(t)* exceeds the firing threshold *V*_*threshold*_, the neuron generates a spike. After firing, *V*_*i*_*(t)* decays exponentially with membrane time constant *τ*_*membrane*_. Let 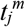 and 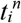 represent the *m*-th and *n*-th spike times of neurons *j* and *i*, respectively.

The synaptic potential *V*_*synapse*_*(t)* depends on the spike timing history of the pre- and postsynaptic neurons. When neuron *j* fires shortly before neuron *i* 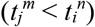, the synapse is potentiated; if the firing order is reversed, the synapse is weakened. The magnitude of synaptic plasticity is modulated by the time difference between spikes, following a spike-timing-dependent plasticity (STDP) rule.

The model also includes an external input term *V*_*external*_*(t)* to account for depolarization due to defined stimuli (e.g., electrical or optogenetic) or stochastic influences (e.g., chemical neuromodulators or glial interactions) [25,26]. Stochastic activity is simulated using a Poisson process, where a randomly selected subset of neurons (*N*_*external*_) generates spikes with interspike intervals (ISI) drawn from an exponential distribution with mean *r*_*external*_. The membrane potential *V*_*i*_ *(t)* evolves according to the leaky integrate-and-fire (LIF) model [24,25,27].

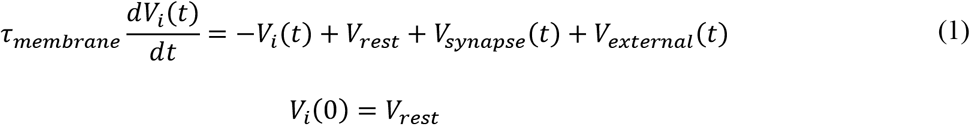

### Synaptic potential, *V*_*synapse*_*(t)*

The synaptic input *V*_*synapse*_*(t)* received by postsynaptic neuron *i* is composed of two terms: a default potential *J*_*ji*_*(t)*, representing baseline vesicle release, and a history-dependent term *s(t)w*_*ji*_ *(t)*, shaped by spike timing and synaptic plasticity. The total synaptic potential is given by:

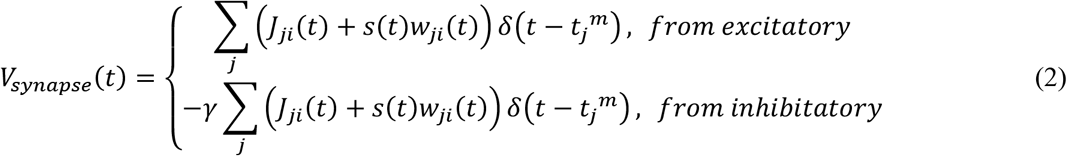

The Dirac delta function δ(*x*) has the property of approaching ∞ when *x = 0*, equaling 0 otherwise, and satisfying the identity 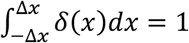 Consequently, when the synaptic potential equation (Eq (1)) is integrated with respect to time from 0 to *t, V*_*synapse*_*(t)* contributes discrete, finite increases to the membrane potential *V*_*i*_*(t)* at each spike time 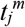 of presynaptic neuron *j* (where *j* = 1, 2, …, and *m* denotes the spike index). The sign of *V*_*synapse*_*(t)* depends on whether the presynaptic neuron is excitatory or inhibitory, determined by the type of neurotransmitter released, typically glutamate for excitatory neurons and GABA for inhibitory neurons [24,25]. The parameter *γ* denotes the ratio of excitatory to inhibitory neurons in the network and influences the overall excitability and balance of the system.

### History-independent component of Vsynapse, J_ji_(t)

The term *J*_*ji*_*(t)* represents the default synaptic potential generated by the release of a baseline number of vesicles. This component is independent of the synaptic firing history but depends on the presynaptic vesicle clustering capacity, which is influenced by mechanical tension. As vesicle accumulation increases under higher tension [10,11,13], we model *J*_*ji*_*(t)* as a function of network-wide mechanical tension (Fig 1C):

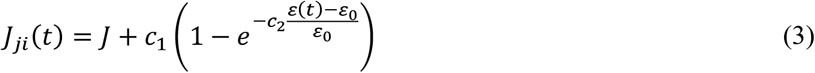

Here, *c*_*1*_, *c*_*2*_, and *J* are constants. The term *J* represents the baseline synaptic potential resulting from vesicle release under a uniform resting tension, denoted as ε_0_, applied across the entire network. When the mechanical tension increases to *ε(t)*, vesicle clustering at the presynaptic terminal is enhanced. This increased vesicle capacity leads to a potentiation of *J*_*ji*_*(t)*, thereby amplifying the synaptic input in a tension-dependent manner.

### History-dependent component of *V*_*synapse*_, *s(t)w*_*ji*_*(t)*

The product *s(t)w*_*ji*_*(t)* represents the plastic, history-dependent component of synaptic transmission. Here, *s(t)* denotes the expected volume of neurotransmitter vesicles released by presynaptic neuron *j* upon firing at time *t*, while *w*_*ji*_*(t)* represents the change in postsynaptic membrane potential per unit volume of released vesicles. Together, this product quantifies the effective contribution to the membrane potential *V*_*i*_*(t)* in response to an action potential from neuron *j*.

### Vesicle release probability, *u(t)*

Let *u(0) = u*_*0*_ denote the baseline probability of vesicle release in the absence of recent synaptic activity (Fig 2A). Upon presynaptic firing, *u(t)* increases due to Ca^2+^ influx at the presynaptic terminal, a well-documented mechanism in neurotransmitter release [28,29]. This increase is transient and decays back toward the baseline *u*_*0*_ as intracellular Ca^2+^ is resorbed, with a time constant *τ*_*u*_. The increment in *u(t)* depends on its value at the moment of presynaptic firing, denoted by 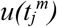, where 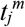 is the *m*-th spike time of neuron *j*. Since the probability is bounded between *u*_*0*_ and 1, the dynamics of *u(t)* is modeled as:

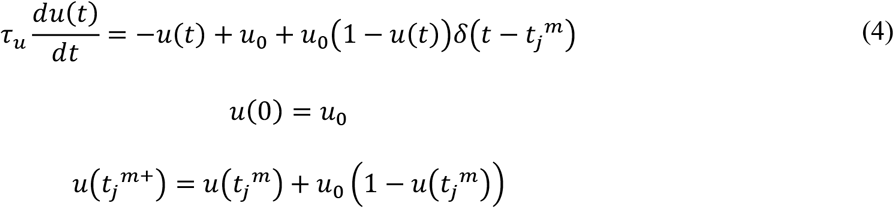

**Fig 2.**
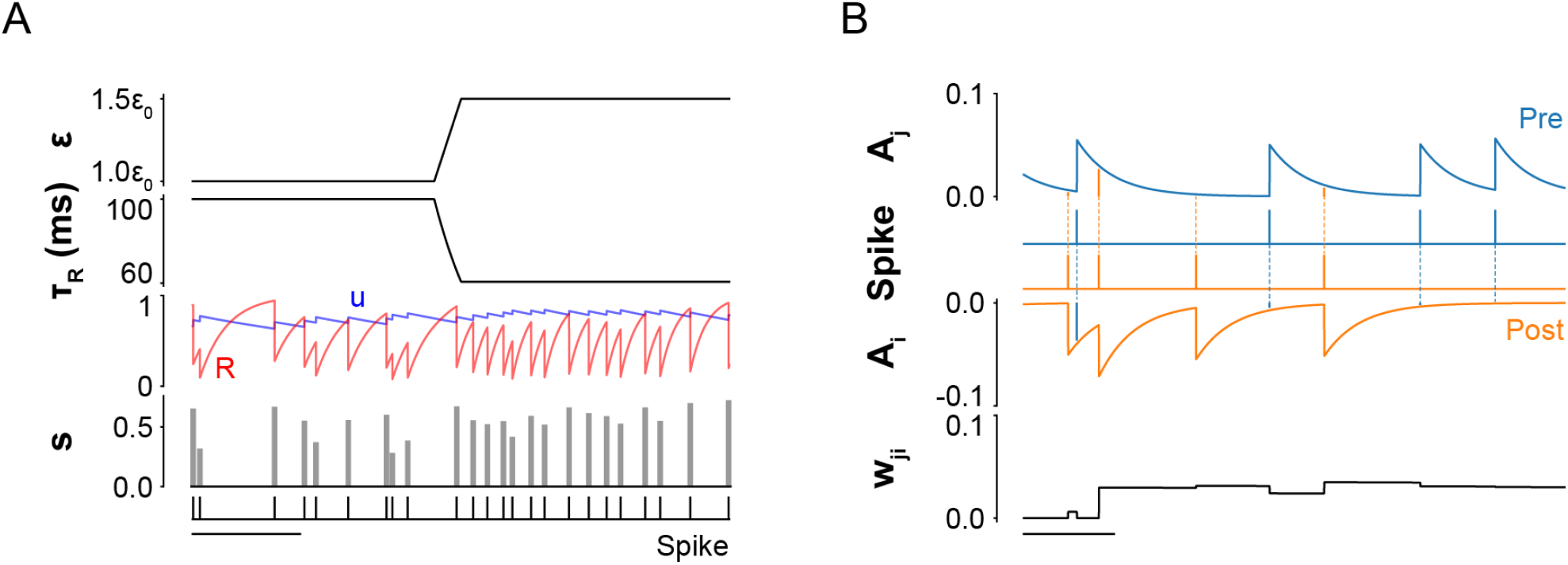
Dynamics of synaptic variables modulated by mechanical tension (A) Vesicle-related dynamics under different tension levels (*ε*). Time course plots show four key variables: vesicle recovery time constant (*τ*_*R*_), release probability (*u*), vesicle availability (*R*), and expected volume of vesicles released (*s*). After a presynaptic neuron fires, *u* increases due to Ca^2+^ influx and then decays exponentially back to its baseline (*u*_*0*_) in the absence of further activity. Simultaneously, *R* decreases as vesicles are released, then recovers toward baseline with time constant *τ*_*R*_. The expected vesicle release volume, *s*, is defined as the product *u* × *R*. Increased tension reduces *τ*_*R*_, thereby accelerating vesicle replenishment and enabling greater vesicle output (*s*) during subsequent spikes(see Materials and Methods: *Table 1*. *All parameters used in simulation* for details of parameters). Scale bar: 400 ms. (B) Time evolution of history-dependent synaptic weight (*w*_*ji*_). Synaptic weight dynamics are shaped by the relative spike timings of the presynaptic neuron *j* and the postsynaptic neuron *i*. Each neuron has a spike trace variable (*A*_*j*_ for the presynaptic and *A*_*i*_ for the postsynaptic neuron) that updates with each spike: *A*_*j*_ increases upon a presynaptic spike and decays back to zero, while *A*_*i*_ decreases upon a postsynaptic spike and also decays to zero over time. *w*_*ji*_ increases by *A*_*j*_*(t*_*i*_) when the postsynaptic neuron fires and decreases by *A*_*i*_*(t*_*j*_) when the presynaptic neuron fires. The magnitude of these changes is modulated by the temporal proximity of the spikes: shorter intervals lead to stronger potentiation or depression. *w*_*ji*_ is bounded between 0 and a maximum value, w_max_. Scale bar: 50 ms.

For instance, if neuron *j* fires for the first time at 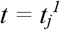, then 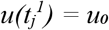, and immediately after firing, 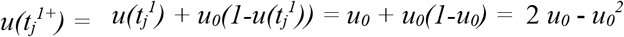. Following this spike-induced increase, *u(t)* decays exponentially back to *u*_*0*_ in the absence of further firing with a time constant of τ_u_.

### Vesicle availability, *R(t)*

When presynaptic neuron *j* fires for the *m*-th time at 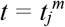, it begins with a default vesicle volume *R*_*0*_ available for release. Immediately after firing at 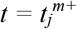, the vesicle pool is depleted proportionally to the release probability, such that 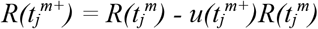. The dynamics of *R(t)* is given by

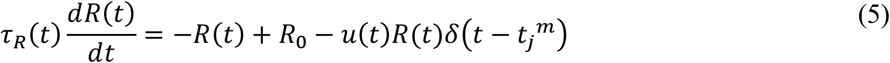

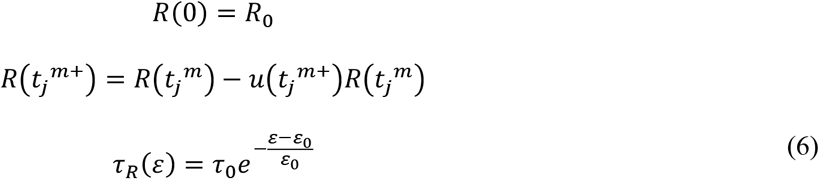

Following this depletion, the vesicle reserve *R(t)* recovers over time with a recovery time constant *τ*_*R*_*(ε)*. This recovery is tension-dependent: as mechanical tension *ε(t)* increases, the recovery becomes faster, that is *τ*_*R*_*(ε)* decreases relative to the baseline *τ*_*0*_.

### Synaptic weight, w_ji_(t)

The synaptic weight *w*_*ji*_*(t)* represents the postsynaptic potential change per unit volume of neurotransmitter vesicles released by presynaptic neuron *j* (Fig 2B). In line with Hebbian theory, we modeled learning-induced plasticity using a STDP framework [26,30,31], which is strongly supported by experimental evidence in hippocampal and cortical neurons [32–35].

Synaptic weights are updated based on the relative timing of spikes from presynaptic neuron *j* and postsynaptic neuron *i*. When *j* fires shortly before 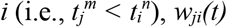 increases, reflecting the strengthening of the synapse. Conversely, when *j* fires after 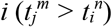, the synapse is weakened, and *w*_*ji*_*(t)* decreases. These updates can be reasoned as follows. When presynaptic firing precedes postsynaptic activation, neurotransmitter binding at the postsynaptic terminal facilitates depolarization and increases synaptic efficacy. In contrast, postsynaptic firing before presynaptic input can trigger back-propagating action potentials and local Ca^2+^ influx through NMDA receptors, leading to attenuation in synaptic potential [32,33,36]. The amount of change in *w*_*ji*_*(t)* depends on the time difference between firing times, 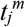 and 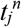, the shorter the time difference, the higher the change.

To implement this in the model, we introduced two temporal traces of pre and postsynaptic neuron: *A*_*j*_*(t)* and *A*_*i*_*(t)*. When neuron *j* fires at 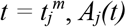 is incremented by a fixed amount *F*_*w*_ and then decays exponentially with a time constant *τ*_*w*_. Similarly, when neuron *i* fires at 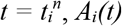 drops sharply and also increases with the same time constant.

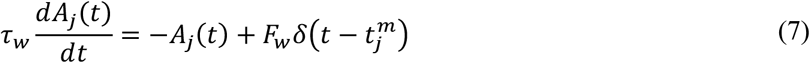

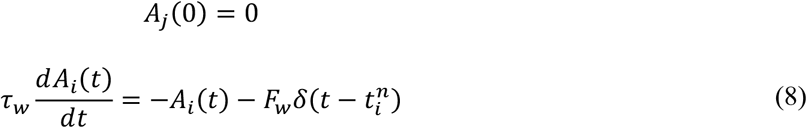

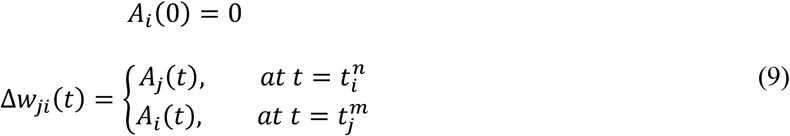

Finally, to prevent unbounded growth, *w*_*ji*_*(t)* decays continuously over time with a forgetting rate *α* and *w*_*ji*_*(t)* is bounded within the range [0, *w*_*max*_].

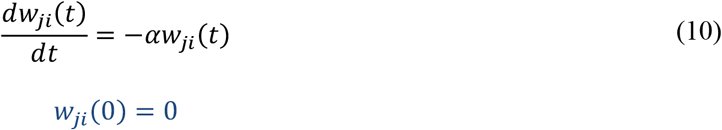

All model parameters are summarized in Table 1.

**Table 1.** All parameters used in simulation.

| Abbreviation | Value | Description | Abbreviation | Value | Description |
| --- | --- | --- | --- | --- | --- |
| $dt$ | 0.1 ms | Simulation time step | $V_{rest}$ | -74 mV | Resting potential |
| $V_{reset}$ | -60 mV | Reset potential | $V_{threshold}$ | -54 mV | Firing threshold |
| $\tau_{membrane}$ | 10 ms | Membrane time constant | | | |
| $N_{external}$ | 1000 | Number of external neurons<br>in the Poisson point process | $r_{external}$ | 13 Hz | Average interval of spikes<br>in the Poisson point process |
| $J$ | 0.01 mV | Vesicle release at resting<br>tension | $\varepsilon_0$ | 0.001 | Resting tension |
| $c_1$ | 0.1 mV | Constant (Eq (3)) | $c_2$ | 0.01 | Constant (Eq (3)) |
| $u_0$ | 0.2 | Vesicle release probability<br>at rest | $\tau_u$ | 1000 ms | Vesicle release probability<br>time constant |
| $R_0$ | 1 | Volume of available<br>vesicles at rest | $\tau_0$ | 100 ms | Vesicle recovery time<br>constant at resting tension |
| $\alpha$ | 0.25 Hz | Forgetting rate | $w_{max}$ | 5 | Maximum synaptic weight |
| $\tau_w$ | 20 ms | Spike trace time constant | $F_w$ | 0.05 | Spike trace amplitude |

### *In silico* stimulation

*In silico* stimulation was performed using a single-pulse stimulation protocol at 50 Hz during training and 10 Hz during recall, applied to predefined regions of the simulated neural network. Mechanical tension was modulated during the recall phase to assess its effect on memory retrieval dynamics. All simulations were conducted in Python (v3.10.13) using standard scientific libraries and previously published neural models [30,37,38].

### Synchrony index between two assemblies

Assembly activity was defined as a period of highly correlated, fast spiking from neurons within a single assembly, characterized by interspike intervals (ISIs) shorter than 0.5 ms. The duration of an assembly activity extended from the onset of the earliest spike to the offset of the latest spike. If an additional neuron fired within 0.5 ms of the most recent spike in the activity, the duration was extended to include that spike. In addition, the instantaneous spike rate at the center of the activity was required to exceed 100 Hz. These criteria were chosen to isolate high-frequency spiking events potentially analogous to ripple oscillations observed *in vivo* and *in vitro*, which are implicated in memory-related neural processes [39–41].

The synchrony index quantified the temporal coordination between two neural assemblies specifically, between a cue-stimulated assembly and another previously co-trained but non–cue-stimulated assembly. To compute the index, assembly activities from both assemblies were vectorized across all detected events. The synchrony index (χ) between two activity vectors, *x* and *y*, was calculated using normalized cross-correlation as follows.

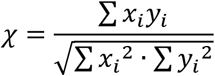

*x*_*i*_ and *y*_*i*_ are the *i*-th elements of the respective vectors. A value of χ = 1 indicates perfect synchrony, while χ = 0 indicates complete asynchrony. Note that *x*_*i*_ or *y*_*i*_ can have values of either 0 or 1.

### Clustering of spiking neurons and intersection over union

To quantify the spatial spread of spiking neurons following stimulation, we applied the density-based spatial clustering of applications with noise (DBSCAN) algorithm [42] over a 5 ms analysis window. DBSCAN identifies clusters by locating core points within dense regions of spiking activity in 2D space and expands each cluster by including neighboring points within a defined radius. A spiking neuron was considered part of a cluster if it met the following DBSCAN parameters:

- Maximum distance between two neurons: 120 μm
- Minimum number of neighboring neurons to form a core point: 20

Once clusters were identified, we computed the spatial similarity between the leaked spiking area (*A*_*leak*_, the area covered by the resulting cluster) and the trained area (*A*_*train*_) using the intersection over union (IoU) metric:

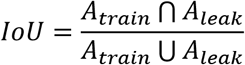

An IoU of 1 indicates perfect spatial overlap between the activated and trained regions, while lower values indicate greater spatial divergence (Fig 3).

**Fig 3.**
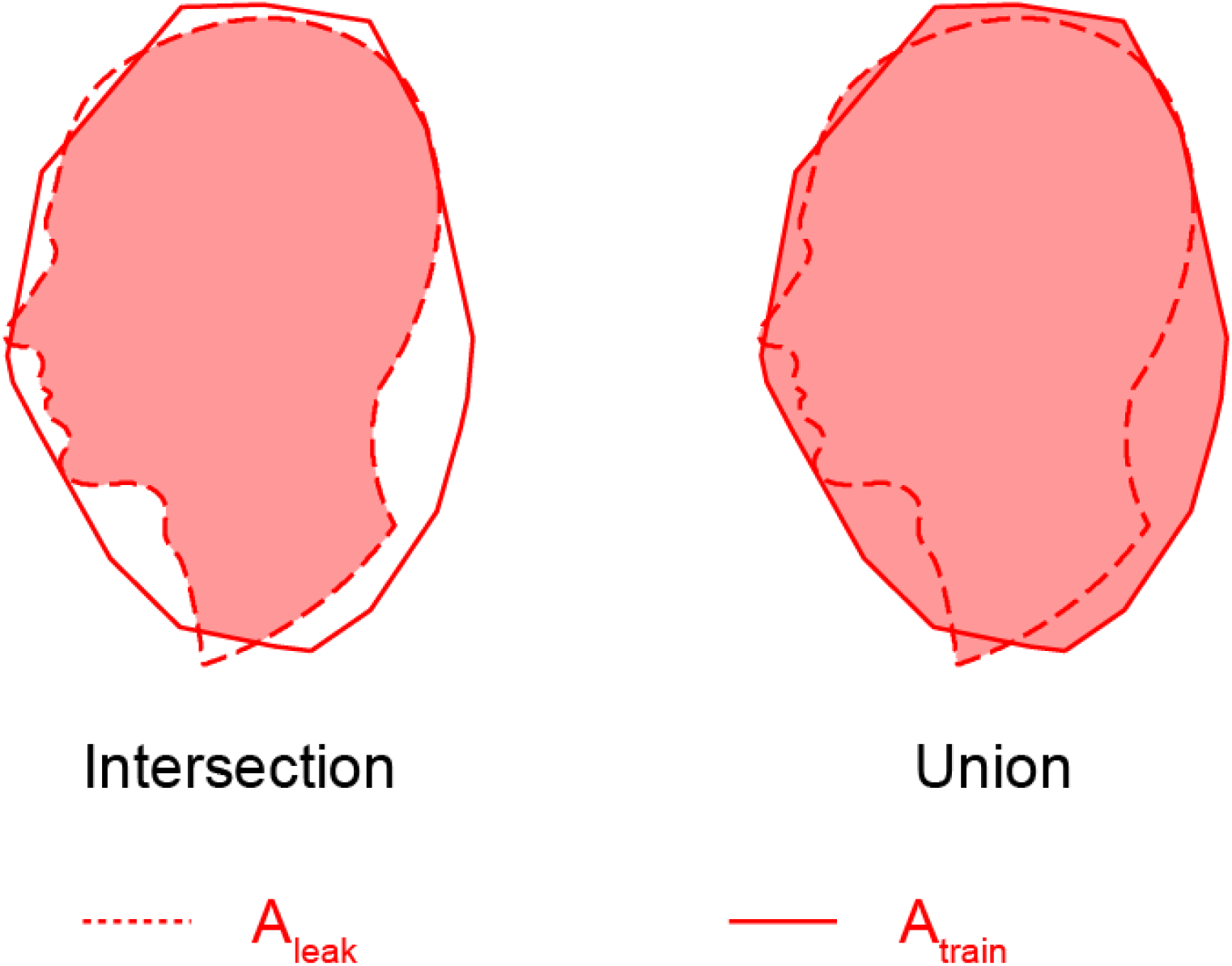
Schematic illustration of intersection over union (IoU). After training the neural network with an image (*A*_*train*_), the spatial distribution of spiking neurons over a 5 ms window was recorded. Upon cue stimulation, the DBSCAN algorithm detected a dense cluster of spiking neurons (*A*_*leak*_). IoU quantifies the spatial similarity between the trained area and the leaked spiking area, calculated as *IoU = A*_*train*_ ∩ *A*_*leak*_*/ A*_*train*_ *∪A*_*leak*_.

### Statistical analysis

Statistical analyses were performed using Python and associated scientific libraries [43]. All data are reported as mean ± standard error of the mean (SEM), with *n* denoting the number of biological or simulated replicates. Statistical significance was set at *p* < 0.05, and significance levels are indicated as follows: *p* < 0.05 (*), *p* < 0.01 (**), and *p* < 0.001 (***). Normality was assessed using skewness and kurtosis thresholds (|skewness| < 1; |kurtosis| < 2), and power transformations were applied to non-normal data to meet these criteria.

Activation time in the pattern completion task and the synchrony index in the projection task were analyzed using one-way ANOVA. Intersection-over-union (IoU) values were evaluated using two-way ANOVA, with the proportion of inhibitory neurons and memory phase (encoding vs. recall) as between-subject factors. Synchrony index in the association task was analyzed using repeated-measures two-way ANOVA, with recall time as the within-subject factor and training duration as the between-subject factor. Post hoc comparisons were conducted using Tukey’s honestly significant difference (HSD) test. F statistics are reported in the form *F*_*m,n*_, where *m* and *n* represent the degrees of freedom between and within groups, respectively.

## Results

### Training duration and tension enhance pattern completion

The network consisted of 4,000 excitatory and 1,000 inhibitory optogenetic neurons, forming both excitatory and inhibitory synapses (Fig 4A). A face-shaped region within the network was selectively trained by exposing light to encode an input pattern. Following training, localized rectangular cues were delivered within this trained region to evaluate whether partial input could elicit recall of the embedded memory via activation of the neural assembly. The total area of the rectangular cues covered 20% of the trained region.

**Fig 4.**
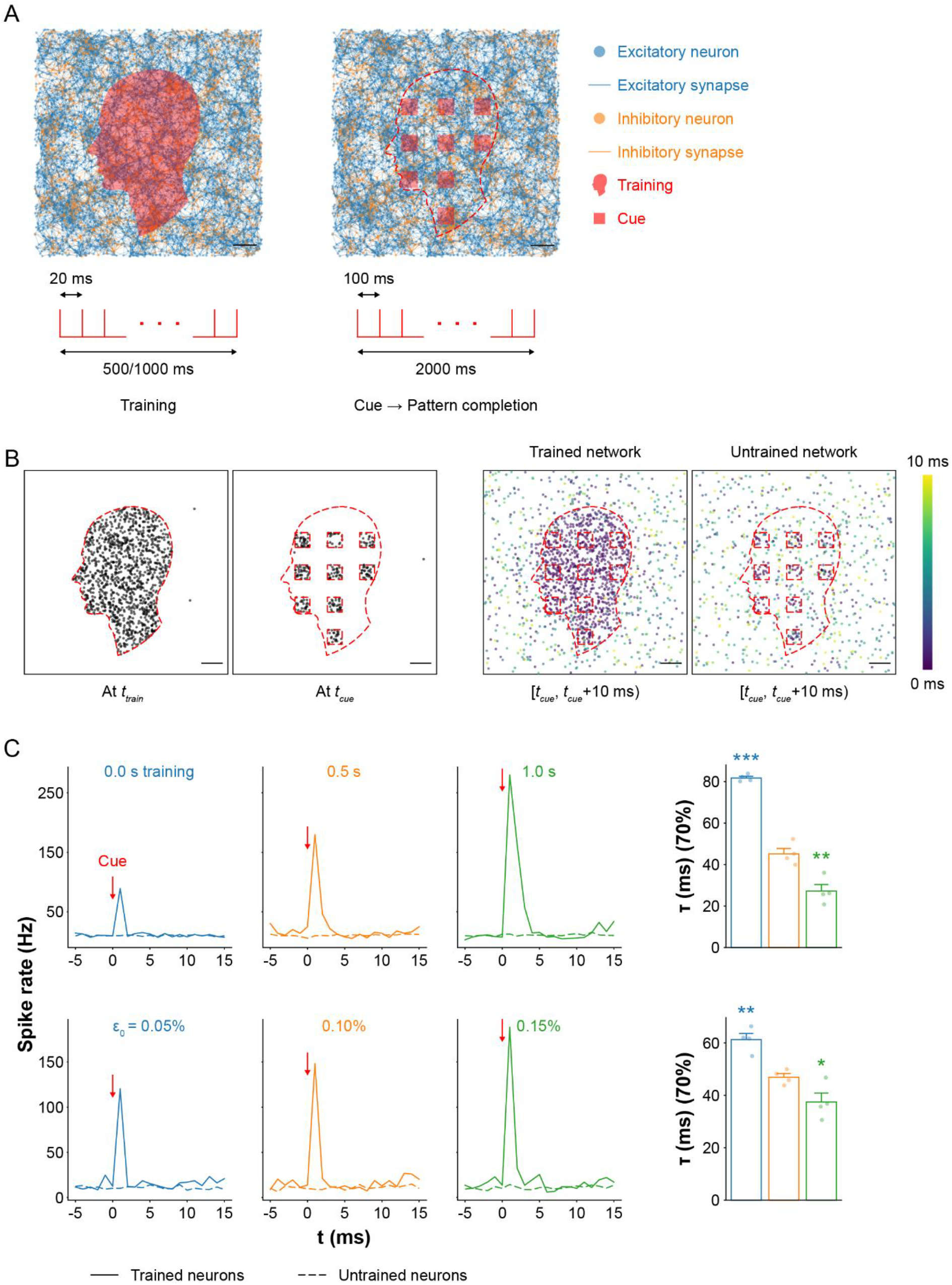
Pattern completion *in silico* under mechanical tension (A) Visualization of spiking neural networks (top) and corresponding stimulation protocols (bottom) during training and recall. During training, neurons within a face-shaped region were repeatedly stimulated. For recall, rectangular subregions covering 20% of the trained area were partially stimulated to assess memory recall. Scale bar: 200 µm. (B) Maps of spiking neurons during training and recall (left), and spiking activity within a 10 ms window following cue stimulation in trained and untrained networks (right). In trained networks, activity propagated from the cue sites across the trained region, while neurons outside the trained area exhibited only spontaneous firing. In contrast, untrained networks showed localized activity restricted to the cue sites, with no propagation. Scale bar: 200 µm. (C) Average spike rates in trained versus untrained regions as a function of training duration (top) and tension level (bottom). Activation times to reach 70% firing in the trained assembly are shown for each condition (right; mean ± SEM; *n* = 4). In the 0.5 s training condition, neurons in the trained region exhibited a sharp spike in activity upon cue stimulation, while neurons in the untrained region showed no change, indicating precise pattern completion. Networks with no training showed a 50% lower peak spike rate, with only a small bump caused by direct activation at the cue site. Longer training (1 s) led to a 56% increase in peak spike rate and sustained firing, along with a 40% reduction in activation time compared to 0.5 s training. Conversely, untrained networks had 81% longer activation times. Increased mechanical tension raised the peak spike rate by 27% and shortened activation time by 20%, while relaxation reduced spike rates by 19% and increased activation time by 31%. Significance levels are shown relative to 0.5 s training or baseline tension (0.1%).

During training, neurons within the face-shaped region responded to the stimuli and fired in a coordinated manner (Fig 4B). In the recall phase, neurons within the cue regions activated in response to the stimuli, and this activity rapidly propagated across the trained area within 10 ms. This spread was driven by the elevated synaptic weights acquired during training. Notably, the activity remained spatially confined to the trained region, as synaptic weights outside this area were insufficient to trigger widespread firing, consistent with localized pattern completion. In contrast, the untrained network showed no propagation beyond the cue site, confirming that training was required for memory-like recall behavior. Dynamic propagation of spikes in both trained and untrained networks is visualized in S1 Video.

To quantify recall performance, we compared spike rates in the trained and untrained regions (Fig 4C). In the 0.5 s training condition, spike rates in the trained region exhibited a sharp increase within 5 ms of cue onset and then decayed rapidly, while spike rates in the untrained region remained low and stable. This indicates that memory-related activity was confined to the trained assembly. The 1 s training condition produced a peak spike rate 56% higher than that of the 0.5 s condition, demonstrating that extended training enhances pattern completion. In contrast, the untrained network showed only a transient spike at the cue site (peak ≈ 89 Hz), followed by rapid decay, reflecting a lack of propagation due to weak synaptic connectivity.

We also measured activation time, defined as the time required for 70% of neurons within the trained region to become active following the cue. Activation time decreased by 40% from the 0.5 s to 1 s training condition (*p* = 0.0049) and increased by 81% in the untrained network (*p* < 0.001; one-way ANOVA, *F*_2,9_ = 129.86, *p* < 0.001), further supporting that training duration significantly enhances the speed of memory recall.

The effect of mechanical tension during recall was also evaluated. Compared to the baseline tension condition, a 50% reduction in network tension resulted in a 19% decrease in spike rate, while a 50% increase in tension enhanced spike rate by 27%. Activation time mirrored these effects: networks under higher tension recalled the pattern 20% faster (*p* = 0.0407), while networks under lower tension recalled 31% more slowly (*p* = 0.0018; one-way ANOVA, *F*_2,9_ = 22.94, *p* < 0.001). These findings suggest that mechanical tension modulates synaptic efficiency, likely through its influence on vesicle availability and recovery dynamics.

Lastly, we investigated whether increasing network size could improve the precision of memory recall, specifically the representation of small facial features such as the nose and lips. To this end, we increased the total number of neurons fourfold to 20,000, resulting in a cell density of 5,000 cells/mm^2^, comparable to previous *in vitro* experiments [9,13] and to estimated neural densities in the human hippocampus, assuming a tissue thickness of 0.25-0.3 mm [44]. The expanded network was trained and cued using the same protocol as before. Spiking activity maps revealed that the 20,000-neuron network reproduced the face pattern better than the 5,000-neuron network (Fig 5), with finer features such as the nose and lips more clearly represented. These results suggest that even a relatively simple model, compared to the highly intricate architecture of the human brain, can exhibit improved memory recall performance with increased neural density.

**Fig 5.**
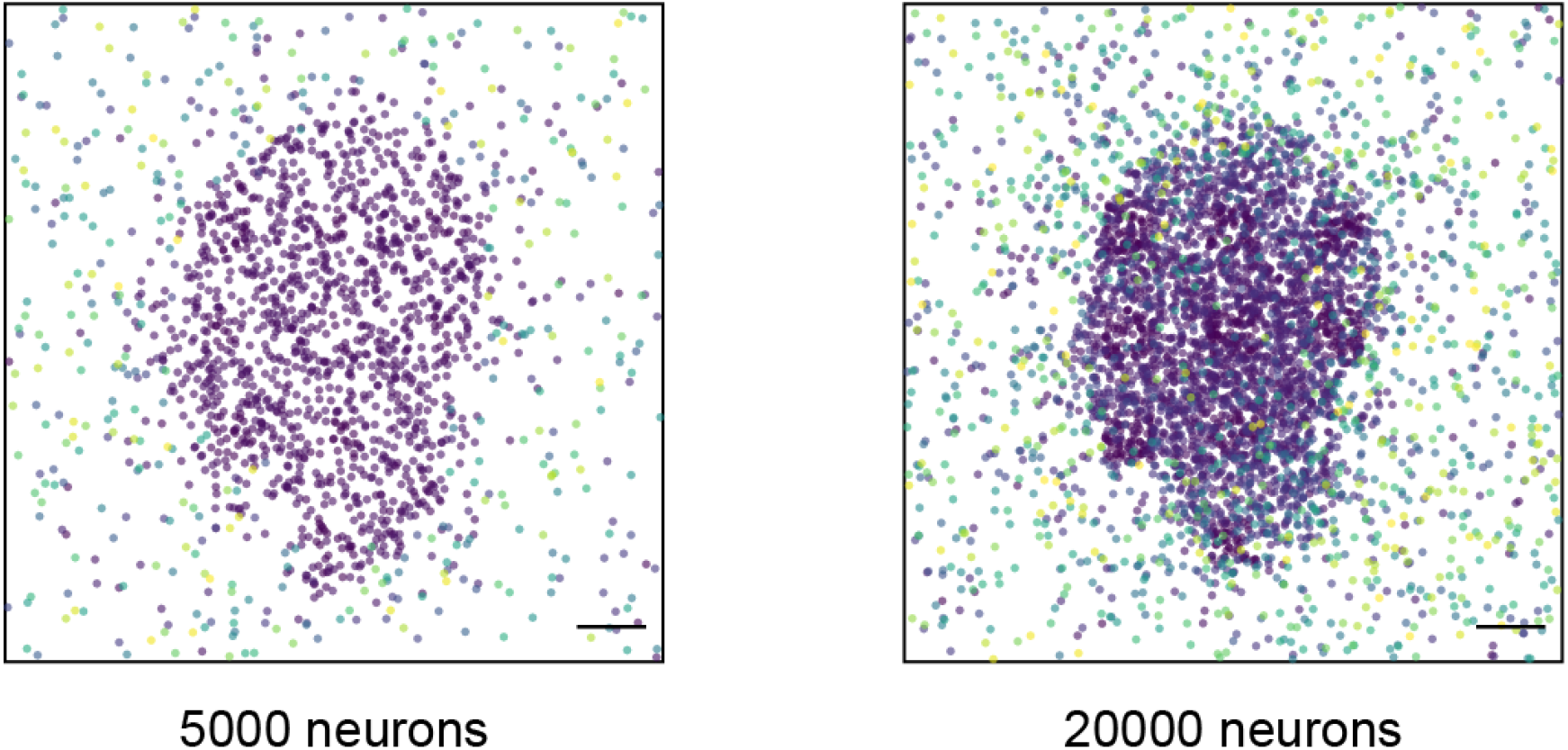
Improvement of memory recall Maps of spiking neurons within a 10 ms window following cue stimulation in network with 5000 (left) and 20000 neurons (right). In network with 20000 neurons, activity propagated more and boundaries of small features such as nose and lips are more visible. Scale bar: 200 µm.

### Inhibitory neurons are critical for memory encoding and recall

To investigate the role of inhibitory neurons in memory formation and retrieval, we constructed networks with varying ratios of inhibitory to excitatory neurons and trained each for 1 second. Memory encoding and recall were assessed by analyzing the spatial spread of spiking activity following training and cue application. Spiking neurons were recorded over a 5 ms window, and cluster size was quantified using DBSCAN (see Materials and Methods: *Clustering of spiking neurons and intersection over union* for details).

During encoding, networks without inhibitory neurons (0%) exhibited widespread spiking beyond the trained region, *A*_*train*_, producing a larger activation area, *A*_*leak*_ (Fig 6A). This resulted from unregulated excitation, as all presynaptic inputs were excitatory. As inhibitory neurons were introduced, *A*_*leak*_ became more confined to *A*_*train*_. In the 100% inhibitory condition, spiking was tightly localized, effectively embedding the stimulus pattern while sharply constraining spread. Thus, in terms of encoding fidelity, networks with more inhibition formed sharper, more well-defined memory traces.

**Fig 6.**
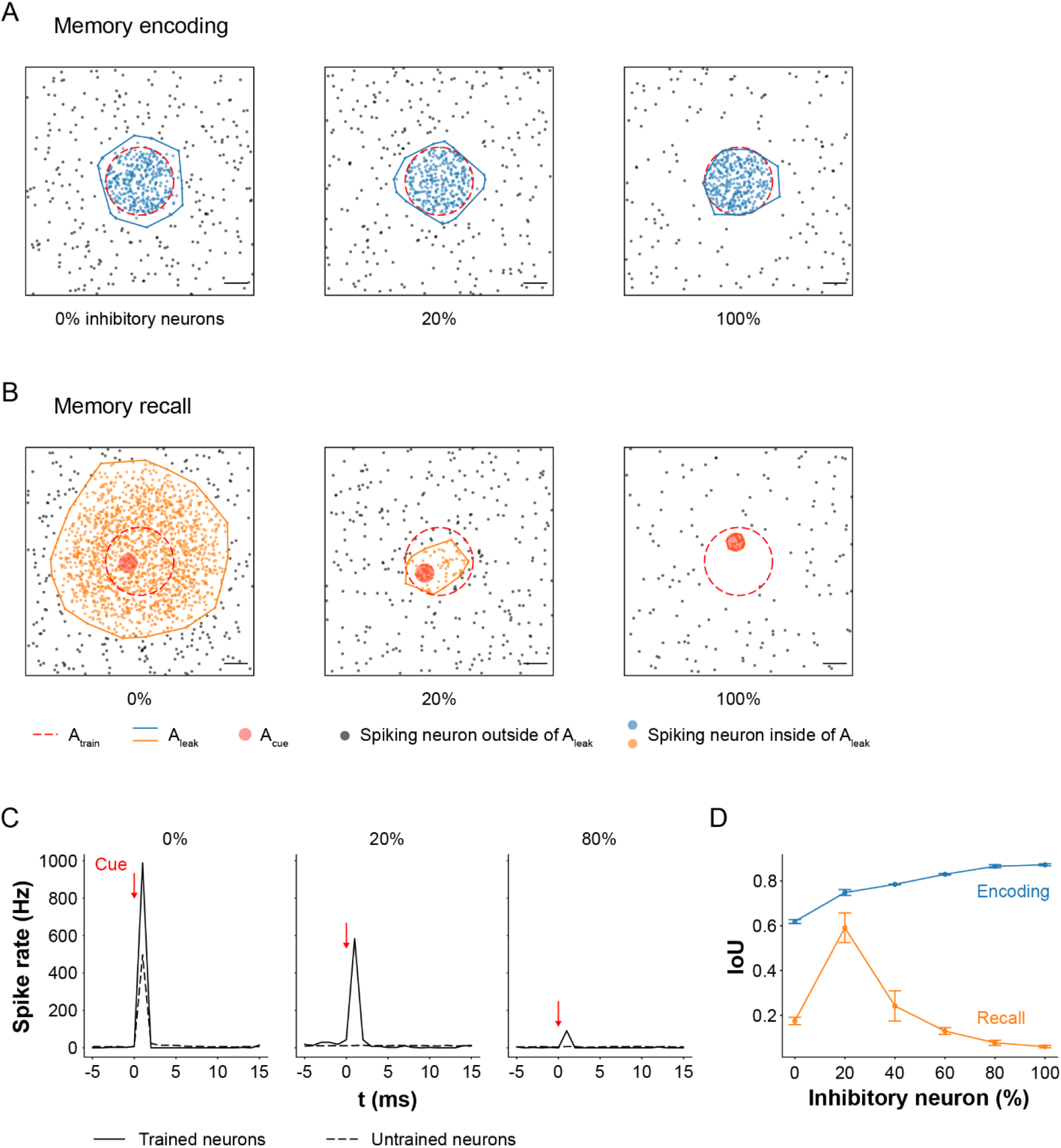
Role of inhibitory neurons in memory encoding and recall (A) Maps of spiking neurons during memory encoding. Clusters of spiking neurons were identified using the DBSCAN clustering algorithm (See Materials and Methods: *Clustering of spiking neurons and intersection over union* for details). In networks with 0% inhibitory neurons, the stimulation-induced spread of activity (*A*_*leak*_) exceeded the trained region (*A*_*train*_) due to unregulated excitatory input increasing the firing probability of postsynaptic neurons. As the proportion of inhibitory neurons increased, *A*_*leak*_ became confined to *A*_*train*_, forming a clearer boundary of the embedded image. Scale bar: 200 µm. (B) Maps of spiking neurons during memory recall in response to partial cue (small circle within the trained region). Networks with 0% inhibitory neurons exhibited uncontrolled propagation of spiking activity from the cue site, indicating imprecise memory recall. At 100% inhibition, activity was limited to the cue site with no propagation, reflecting recall failure. In contrast, networks with 20% inhibitory neurons showed more localized and accurate recall of the trained image. Scale bar: 200 µm. (C) Spike rates during recall for networks with 0%, 20%, and 100% inhibitory neurons. Networks composed entirely of excitatory neurons (0%) showed the highest spike rates due to lack of inhibitory control. Conversely, networks with all inhibitory neurons (100%) showed minimal activity. Networks with 20% inhibition achieved moderate but targeted firing, consistent with effective recall. (D) Intersection over union (IoU) values between *A*_*leak*_ and *A*_*train*_ across inhibitory neuron ratios during encoding and recall (*n* = 4). At 0%, the lowest IoU during encoding reflected poor memory formation, which resulted in poor recall. Introducing inhibitory neurons improved IoUs, with the highest encoding IoU at 100%, but this dropped by 93% during recall, indicating over-suppression. Optimal memory retrieval occurred at 20% inhibition, where recall IoU was highest. Although recall IoU was 21% lower than encoding IoU at this ratio, the difference was not statistically significant.

However, recall revealed a different pattern. In 0% inhibitory networks, the cue-induced activity spread uncontrollably, resulting in diffuse and imprecise activation that extended far beyond *A*_*train*_ (Fig 6B). In contrast, 100% inhibitory networks showed minimal activity beyond the cue site, suggesting a failure to retrieve the full embedded pattern despite successful encoding. Notably, networks with 20% inhibitory neurons achieved both containment and propagation of activity, accurately recalling the stored pattern with high spatial precision. These patterns were also reflected in spike rate measurements during recall (Fig 6C). Networks without inhibition (0%) showed the highest spike rates, reflecting runaway excitation. In contrast, fully inhibitory (100%) networks showed a minimal increase in spike rate, limited to the immediate cue region. The 20% condition showed an intermediate but structured spike rate response, consistent with controlled and targeted recall.

To quantify memory fidelity, we computed IoU between *A*_*train*_ and *A*_*leak*_. Statistical analysis revealed significant effects of inhibitory ratio (*F*_5,36_ = 20.29, *p* < 0.001), phase (encoding vs. recall; *F*_1,36_ = 1194.73, *p* < 0.001), and their interaction (*F*_5,36_ = 37.45, *p* < 0.001). Overall, IoUs during encoding were significantly higher than during recall (*p* < 0.001), reflecting the probabilistic nature of recall i.e., retrieval does not perfectly reproduce the original activation pattern on every trial.

During encoding, the 0% inhibitory condition yielded the lowest IoU, indicating the poorest encoding quality. IoU increased with higher inhibitory neuron ratios, reflecting more spatially confined memory formation. During recall, the lowest IoU (0.1748) was observed in the 0% condition, due to poor encoding and uncontrolled spread of activity. In the 100% inhibitory case, the IoU difference between encoding and recall was largest, decreasing by 93% (*p* < 0.001) suggesting strong encoding but near-total failure of recall.

The optimal recall performance occurred at a 20% inhibitory neuron ratio, which yielded the highest recall IoU. At this ratio, the recall IoU was only 21% lower than the encoding IoU (0.7900-fold), and the difference was not statistically significant (*p* = 0.0586), indicating that the network was able to recall approximately 79% of the actual image embedded during training. These findings support the hypothesis that an appropriate balance of excitation and inhibition is essential for both precise memory encoding and successful recall.

### Tension enhances projection between assemblies

Projection, an advanced cognitive operation, refers to the activation of a downstream assembly (B) by an upstream assembly (A), effectively “projecting” learned information across regions. To model this process, we trained two spatially separated assemblies within the network (Fig 7A). During training, region A (square) received full stimulation, while region B (triangle) was partially stimulated 2 ms later (40% of neurons), to encourage directed connectivity from A to B without forming an independent assembly in B.

**Fig 7.**
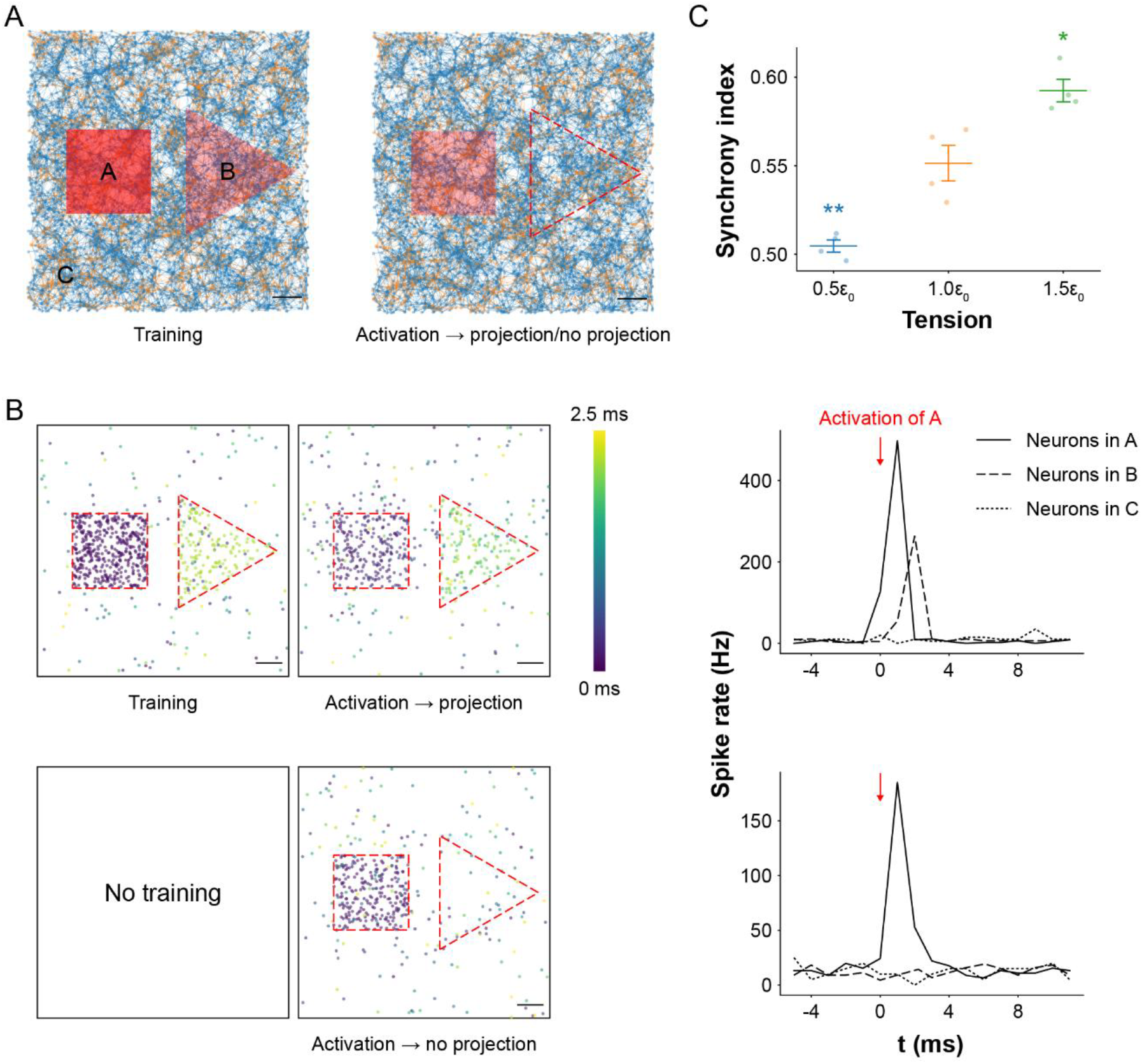
Projection between neural assemblies *in silico* (A) Visualization of neural networks during training and projection. The transparency of stimulated regions represents relative stimulation intensity. During training, neurons in region A were fully stimulated (same protocol as in pattern completion) to form a neural assembly. Neurons in region B were stimulated 2 ms after region A and at 40% of A’s stimulation intensity to encourage the formation of connectivity from A to B without creating an independent assembly in B. Scale bar: 200 µm. (B) Maps of spiking neurons within a 2.5 ms window after activation of region A during projection, comparing trained (top left) and untrained (bottom left) networks. Corresponding spike rates in regions A, B, and C are shown during projection for trained (top right) and untrained (bottom right) conditions. In the trained network, neurons in region A responded strongly during training, while region B showed delayed and weaker activation due to partial stimulation. During projection, activation of A induced activity in B with a short lag, indicating successful projection. Region C exhibited no activity, confirming that it is not involved in the projection. In contrast, the untrained network showed no projection from A to B, as reflected by the absence of spiking activity in B. Scale bar: 200 µm. (C) Synchrony indices between assemblies A and B under different tension levels (mean ± SEM; *n* = 4). Higher tension (150% of baseline, i.e., 1.5ε_0_) significantly increased synchrony by 7%, while lower tension (50% of baseline, i.e., 0.5ε_0_) decreased synchrony by 8%, relative to the resting tension (1.0ε_0_). These results indicate that mechanical tension enhances projection between neural assemblies. Significance levels are shown with respect to the baseline condition.

During training, most neurons in region A responded and fired in response to stimulation, whereas a smaller subset of neurons in B fired with a 2 ms delay due to the weaker, delayed input (Fig 7B). After training, activation of region A alone was sufficient to trigger delayed firing in region B, indicating successful projection from A to B. In contrast, no such projection was observed in the untrained network, where activity in A failed to elicit any response in B. This dynamic propagation is visualized in S1 Video.

Spike rate profiles further support this observation. In the trained network, activation of A led to a spike rate surge in region B with a consistent short delay (∼2 ms). No changes were observed in an unrelated control region (C), confirming that projection was specific to the A→B connection. In the untrained network, spike rates in A were lower, reflecting weaker intra-assembly connections, and no activity was detected in B, confirming the absence of projection.

To quantify projection strength, we computed the synchrony index between assemblies A and B (see Materials and Methods: *Synchrony index between two assemblies* for details). Mechanical tension significantly modulated projection efficacy (one-way ANOVA *F*_2,9_ = 38.05, *p* < 0.001; Fig 7C). Increasing tension from the resting level (100% to 150%) enhanced the synchrony index by 7% relative to baseline (*p* = 0.0134), suggesting that tension facilitates inter-assembly communication. Conversely, reducing tension to 50% of resting levels led to an 8% decrease in synchrony (*p* = 0.0044), likely due to decreased vesicle availability and impaired synaptic transmission. Together, these results suggest that mechanical tension not only affects individual neuron firing but also governs the functional coupling between assemblies necessary for information flow.

### Tension and training duration modulate associative memory recall

We investigated associative memory, a higher-order cognitive function involving the functional linking of distinct neural assemblies. To model this process, we simultaneously trained two spatially separated rectangular regions (see Materials and Methods: *In silico stimulation* for details). To test associative recall, we applied a partial cue by stimulating only the right rectangular region and monitored activity in the unstimulated left region (Fig 8A). We also manipulated mechanical tension to examine its influence on memory accessibility.

**Fig 8.**
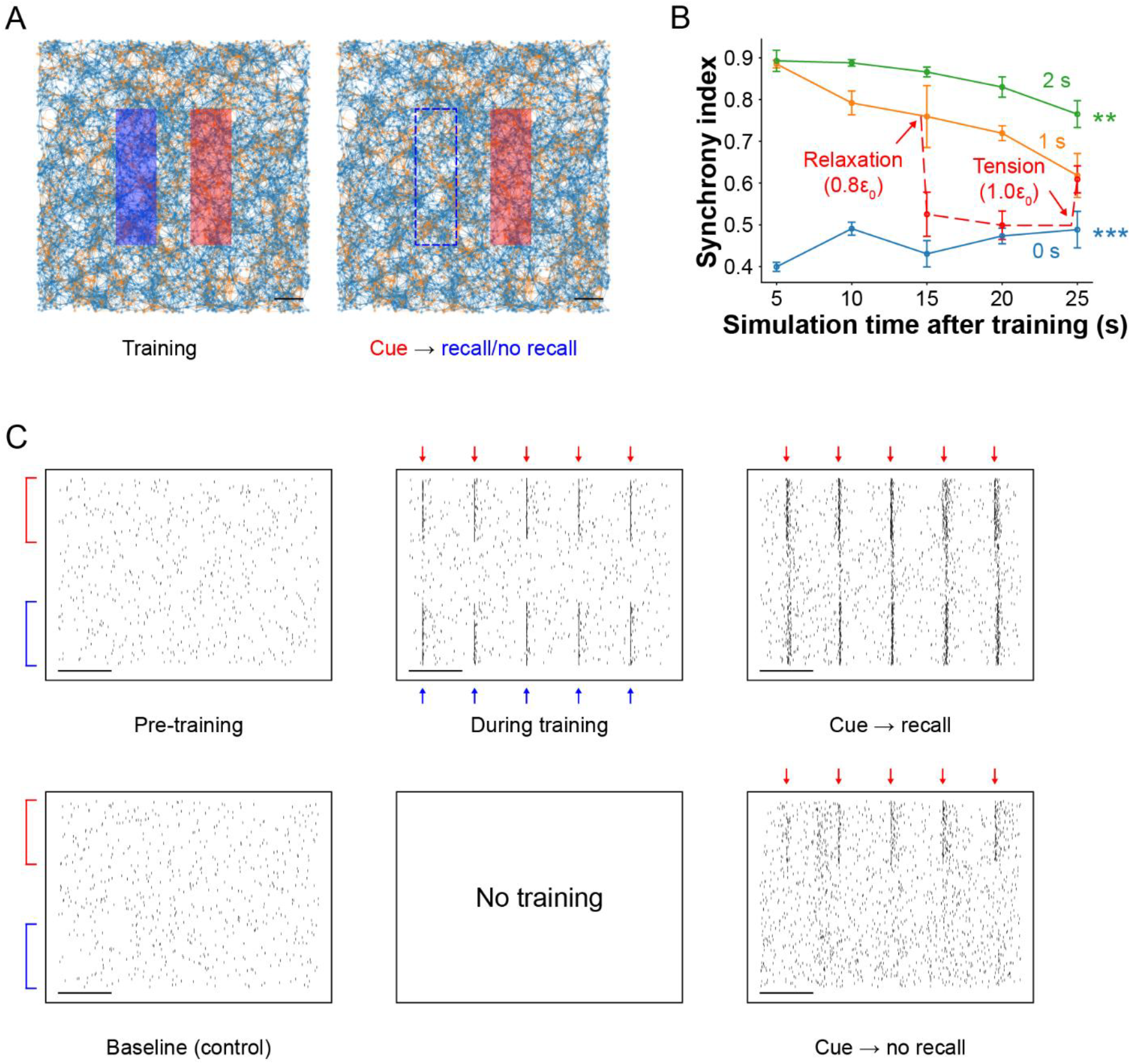
Memory association *in silico* (A) Visualization of the spiking neural network with two spatially separated stimulated regions. During training, both the red and blue regions were simultaneously stimulated. During recall, only the red region received cue stimulation (same protocol as in pattern completion). Scale bar: 200 µm. (B) Synchrony indices between the two regions during cue stimulation following 0, 1, and 2 seconds of training (mean ± SEM; *n* = 4). The model exhibited memory-like dynamics, including memory formation, cue-induced recall, and decay. Networks trained for 1 s showed higher synchrony than untrained networks, though synchrony declined over time, reflecting memory fading. Two seconds of training resulted in significantly higher and more persistent synchrony, indicating stronger memory formation. Mechanical tension also modulated recall: reducing tension by 20% led to a 31% drop in synchrony, which remained suppressed at 20 seconds, indicating impaired memory access. However, restoring tension to baseline recovered synchrony to its original level, suggesting that the memory trace remained intact but was temporarily inaccessible under reduced tension. Significance levels are reported relative to the 1-second training condition. (C) Raster plots of spiking activity at baseline, during training, and during recall in trained (top) and untrained (bottom) networks. The y-axis shows consistent electrode indices across conditions. Red and blue arrows indicate stimulus application. In trained networks, cue stimulation during recall triggered synchronous firing in the previously co-stimulated blue region, demonstrating memory association. Untrained networks exhibited no such synchrony in the blue region. Scale bar: 20 ms.

During training, neurons across both regions exhibited temporally coordinated spiking, reflecting their joint activation by the composite stimulation pattern (Fig 8C). In the recall phase, stimulation of the right region alone evoked synchronous bursts not only locally but also in the left region, indicating functional linkage and associative memory recall. In contrast, when the same cue was applied to an untrained network, only spontaneous, uncoordinated activity appeared in the left region, indicating no stored association.

We quantified recall performance using the synchrony index between the two assemblies (See Materials and Methods: *Synchrony index between two assemblies* for details; Fig 8B). A repeated-measures two-way ANOVA revealed significant effects of recall time (*F*_4,36_ = 5.84, *p* < 0.001), training duration (*F*_2,9_ = 131.23, *p* < 0.001), and their interaction (*F*_8,36_ = 4.87, *p* < 0.001). The trained networks exhibited high synchrony during training, driven by co-stimulation, and maintained elevated synchrony during recall compared to untrained controls (*p* < 0.001). However, synchrony declined gradually over 25 s of recall, indicating memory decay. Networks trained for 2 s showed significantly higher synchrony than those trained for 1 s (*p* = 0.0034), suggesting that longer training enhances both strength and longevity of memory associations.

To evaluate the role of tension, we reduced network tension by 20% following 1 s of training. This resulted in a 31% drop in synchrony index (*p* = 0.0423), which remained significantly suppressed after 20 s (*p* = 0.0012). Crucially, restoring tension to baseline fully recovered synchrony levels (*p* = 0.8863), indicating reinstatement of associative recall. These findings suggest that memory was not erased during relaxation but rendered temporarily inaccessible, likely due to reduced vesicle availability and impaired synaptic efficacy. Restoring tension reinstated effective synaptic transmission and re-enabled memory retrieval.

## Discussion

In this study, we developed an *in silico* platform to investigate how the mechanical tensional state of neural networks modulates high-level cognitive operations, including learning, memory, projection, and association. While recent studies have shown that mechanical tension can influence synaptic plasticity, vesicle accumulation, and neural excitability [3–5,9–13], our work advances this line of research by integrating tension directly into a biologically inspired spiking neural network model. The model predicts that increased tension enhances synaptic function by promoting vesicle clustering at presynaptic terminals and accelerating vesicle replenishment. These tension-driven improvements in synaptic dynamics translate into faster memory recall, stronger neural synchrony, and more reliable execution of cognitive operations, suggesting that mechanical tension may serve as a key modulator in both biological and artificial neural systems.

We modeled learning-based plasticity using spike-timing-dependent plasticity (STDP), allowing synaptic weights to adapt based on the temporal order of pre- and postsynaptic spikes. This mechanism enabled the network to encode and recall learned information. However, to more faithfully capture the dynamics of brain activity, future models should incorporate neural bursting. Bursting, defined as consecutive spikes recorded from a single electrode *in vivo* or *in vitro*, has been linked to memory-related processes and can arise from multiple mechanisms, including the spatial organization of synaptic inputs [45], ionic dynamics involving voltage-gated calcium channels [46], and modulation of Ca^2+^ conductance via G protein-coupled receptors such as metabotropic glutamate receptors (mGluRs) [47]. In humans, ripple oscillations, transient high-frequency bursts exceeding 80 Hz, are thought to support memory encoding and recall [39–41,48]. However, our computational model did not reproduce bursts at the single-neuron level. Given the functional significance of bursting in cognitive operations, future models should incorporate burst-generating mechanisms to more accurately capture how mechanical tension may shape biologically realistic patterns of memory and learning.

A key insight from our model was the essential role of inhibitory neurons in memory encoding and recall. In networks lacking inhibition (0% inhibitory neurons), memory encoding was poor due to the unbounded excitatory activity of presynaptic neurons, and cue stimulation triggered uncontrolled propagation of spikes far beyond the trained region. This is consistent with the well-established role of inhibition in maintaining network stability and regulating sensory-evoked responses [49,50]. Conversely, in networks composed entirely of inhibitory neurons (100%), we observed the formation of clearly defined memory traces during training due to suppressed firing at the boundary of the trained region but during recall, activity remained strictly confined to the cue site and failed to propagate, resulting in poor memory retrieval. The optimal encoding and recall occurred at an intermediate configuration, approximately 20% inhibitory neurons, where excitation was sufficiently constrained to allow precise and localized memory recall. This result is broadly consistent with anatomical data: in mice and rats, 7–11% of hippocampal neurons are GABAergic interneurons (inhibitory), with the remainder being excitatory [20,21]. In primates, GABAergic neurons constitute approximately 20–25% of cortical neurons [22,23]. Together, these findings reinforce the idea that inhibitory neurons are not merely suppressive but play a critical regulatory role, refining and stabilizing neural representations to support accurate memory encoding and recall while preventing pathological excitation and network instability.

Our findings on pattern completion, projection, and association invite a deeper reinterpretation of assembly calculus in biological terms. Assembly calculus conceptualizes neural assemblies as symbolic representations of concepts such as images or words, and models cognitive operations using abstract logical rules (e.g., “if A fires, then B fires”) [17]. However, our work demonstrates that such high-level operations can emerge from low-level, biologically plausible mechanisms, including STDP and vesicle dynamics. Notably, we observed that the success of projection and association was modulated by mechanical tension, suggesting the presence of a neuromechanical layer in cognitive control.

This raises the possibility that tension acts not only as a modulator of firing and plasticity, but as a control mechanism, determining when, where and what conditions assemblies can interact with each other. In this view, mechanical forces impose real-world, physical constraints on otherwise abstract symbolic operations, effectively bridging the gap between continuous biological dynamics and discrete computational logic. Taken together, our results suggest that high-level cognitive functions may emerge not solely from fixed logical architectures, but from the interplay of structural connectivity, plasticity, and physical state, pointing toward a more integrated and dynamic model of cognition.

Another compelling implication of our model lies in its potential relevance to clinical memory disorders such as Alzheimer’s disease, dementia, and amnesia. In our simulations, relaxing mechanical tension in the network led to a significant drop in the synchrony index, indicating impaired memory recall. However, when tension was restored to its original level, synchrony returned to baseline suggesting that the memory trace itself had not been erased, but had become temporarily inaccessible under low-tension conditions. This finding aligns with prevailing hypotheses in clinical neuroscience, which propose that patients with memory impairments may not suffer from the loss of stored memories, but rather from disrupted access to intact memory traces [51,52].

Mechanistically, our model offers a plausible explanation: reduced tension leads to diminished vesicle clustering and slower vesicle replenishment, impairing synaptic transmission and preventing the network from reactivating previously learned assemblies, without compromising synaptic efficacy that was acquired through synaptic plasticity. When tension is reinstated, vesicle availability is restored, allowing memory recall to resume due to synaptic efficacy acquired during learning before reduction of tension. These results suggest that the mechanical state of neural tissue may play a direct and reversible role in determining whether memories can be accessed, rather than whether they are stored at all, providing a new interpretive lens for memory systems.

In summary, we demonstrated that mechanical tension enhances key cognitive operations, including pattern completion, projection, learning, and memory, within a biologically inspired spiking neural network model. These findings provide a conceptual foundation for the development of biocompatible or biohybrid computing systems that exploit physical principles such as tension for learning and memory. More broadly, they advance our understanding of mechanical tension as an active neuromodulatory factor in cognitive function. Future work can build on this framework to investigate how mechanical states influence other brain functions, including computation, reasoning, planning, language, and forgetting, opening new directions across neuroscience, artificial intelligence, and neuroengineering.

## Supporting information captions

S1 Video. Pattern completion with and without training.

Spatiotemporal propagation of spiking activity upon cue stimulation in trained (left) and untrained (right) networks. In the trained network, spiking initiated at the cue sites rapidly propagates across the trained region, demonstrating successful pattern completion. In contrast, the untrained network shows restricted activity confined to the cue sites, with no propagation. Each frame represents neural activity within a 1 ms time window. Total duration: 15 ms. Scale bar: 200 μm.

S2 Video. Projection with and without training.

Spatiotemporal propagation of spiking activity following activation of assembly A in trained (left) and untrained (right) networks. In the trained network, activation of A successfully triggers projection to assembly B with a short temporal lag. In contrast, the untrained network shows no projection from A to B. Each frame represents neural activity within a 1 ms time window. Total duration: 15 ms. Scale bar: 200 μm.

